# Quantitative proteomics reveals the protective effects of ESD against osteoarthritis via attenuating inflammation and modulating immune response

**DOI:** 10.1101/2020.07.15.204552

**Authors:** Ying Hao, Yang Wu, Shanglong Wang, Chungguo Wang, Sihao Qu, Li Li, Guohua Yu, Zimin Liu, Zhen Zhao, Pengcheng Fan, Zengliang Zhang, Yuanyuan Shi

**Author notes:** These authors contributed equally to this work as co-first authors (Ying Hao, Yang Wu, Shanglong Wang). Correspondence to: Yuanyuan Shi,; Zengliang Zhang,; Pengcheng Fan,.

## Abstract

*Epimedium, Salvia miltiorrhiza*, and *Dioscorea nipponica* Makino (ESD) have been combined to treat osteoarthritis (OA) for a long time. In this study we used quantitative proteomics to find the protective effects of ESD against OA and possible mechanism. After papain-induced rats’ OA model established ESD was intragastrically administrated to rats for four weeks. Label-free quantitative proteomics was used to screen the comprehensive protein profiling changes in both OA and ESD groups. After stringent filtering, 62 proteins were found to be significantly up-regulated and 208 proteins were down-regulated in OA model compared with sham-operated control. Functional analysis revealed that these OA up-regulated proteins were enriched in the activation of humoral immunity response, complement activation, leukocyte mediated immunity, acute inflammatory, endocytosis regulation, and proteolysis regulation. ESD partially recovered the protein profiling changes in OA model. The effects of ESD were also assessed by measurement of behavioral activity and pathologic changes in the joints. ESD showed protective effects in suppressing inflammation, releasing joint pain, and attenuating cartilage degradation. Our study presented that ESD as a potential candidate to alleviate OA damage by reducing inflammation and modulating of immune system.

---

Osteoarthritis (OA) is the most common form of arthritis in the world and has a major effect on the health-related quality of life (1). It has long been viewed as a degenerative disease caused by insufficient regeneration of cartilage in joints, most often in the fingers, knees and hips (2, 3). Common clinical features include joint pain, difficulties walking and disability (4, 5). Although glucosamine and chondroitin sulfate have been used widely as dietary supplements for OA (6). However, the effectiveness of these supplements remains controversial (6–8). The efficacy of nonsteroidal anti-inflammatory drugs was challenged with safety and tolerability issues (7). As the present drugs’ limitations, more effective therapeutic strategies are needed to prevent OA progression.

Natural herbal agents have been widely used in clinical application for OA treatments (9, 10). *Epimedium, Salvia miltiorrhiza* and *Dioscorea nipponica* Makino (ESD) as a compound preparation has been proved to be effective in clinical OA therapy (11–16). However, the protein profiling protective effects of ESD were still unclear.

Mass spectrometry (MS) based proteomics provided feasible tools for insights into the patterns of protein expression (17, 18). Here, we used label-free quantitative approaches to find proteomics protective effects of ESD against OA and potential mechanisms. A stable rat OA model was established to evaluate the pharmacological effects of EDS.

## EXPERIMENTAL PROCEDURES

*Preparation of ESD-The* ESD (JointAlive^™^, Chenland Nutritionals, Inc., California) was prepared by mixing *Epimedium, Salvia miltiorrhiza*, and *Rhizoma Dioscoreae Nipponicae* powder with the ratio of 360:24:216 (*w*/*w*).

### LC-MS Analysis of ESD Solution

The methanol and acetonitrile used for the mobile phase (HPLC grade) and reagents used for the sample preparation (analytical grade) were obtained from Merck (Darmstadt, Germany). ESD powder (1 g) was dissolved in 10 mL of methanol, and extracted by ultrasonic for 60min. After filtering the extracts, 1 mL of the filtrate was mixed with 9 mL methanol. 1 mL of the diluted solution was centrifuged at 12000 rpm for 10 min. The supernatant was filtered through 2.2 μm filter membrane for subsequent mass spectrometry analysis. Q Exactive Plus mass spectrometers (Thermo Fisher Scientific, Rockford, IL, USA) coupled with an Ultimate 3000 UPLC system (Thermo Fisher Scientific) was used for the component analysis. The column temperature was 30 °C. The injection volume was 3 μl. The flow rate was 0.3 mL·min^-1^ with mobile phase ACN (0.1% Formic acid) from 5% to 80% for 25 minutes. The LC-MS/MS scan range was from 50 to 1500 m/z. The components were identified by m/z searching in the Traditional Chinese Medicine Systems Pharmacology Database and Analysis Platform (TCMSP).

### Rat Knee OA Model and Treatment

All Wistar rats (280±30 g, half male and half female) used in this experiment were SPF animals and obtained from Qing Longshan Experimental Animal Center (Nanjing, China). The study was approved by the Committee on the Ethics of Animal Experiments of Beijing University of Chinese Medicine. Animals were allowed access to food and water *ad libitum*. The animals were kept on a 12-h day-night cycle. The animals were subjected to adaptive feeding for 7 days prior to initiation of the experiments. Sixty rats were divided into five groups (*n*=12): control group; OA group (OA group); glucosamine and chondroitin Sulfate group (GA group); *Epimedium brevicornu, Salvia miltiorrhiza, Dioscorea nipponica* Makino Low-dose group (ESDL group) and *Epimedium brevicornu*, *Salvia miltiorrhiza*, *Dioscorea nipponica* Makino High-dose group (ESDH group). The OA model was prepared according to the reference (19, 20). Rats in the control group were injected with 0.1 mL of saline and rats in the OA groups were injected with 0.1mL 4% papain solution on the first day of the experiment. The injection position was at the outer edge of the inferior patellar tendon to the intermalleolar fossa. The same treatment was carried out on day 1st, 4th, and 7th respectively, and 4% papain solution was injected three times in total. Each rat was administrated by gavage once a day from the day of modeling. In the placebo group, the same volume normal saline was administrated every day until 28 days.

### Protein Extraction, Digestion and Enrichment

In each group twelve samples were pooling into three, then following protein extraction and trypsin digestion. We homogenized 20 mg pieces of frozen mouse articular cartilage of knee in 0.20 mL of 0.1 M Tris-HCl, pH 7.6 using a rapid low temperature tissue homogenizer (Tissuelyser®, Jingxin Industrial Development Co., Ltd., Shanghai, China) at power 60 Hz at −20 °C for 120 s. The homogenate was centrifuged at 13,300g at 4 °C for 10 min. The supernatant protein concentration was determined by the protein quantification kit (Dingguo Changsheng, Beijing, China) according to the instructions. The protein solution samples were followed by process of filter-aided sample preparation (FASP) (21). Briefly, Aliquots of lysates corresponding to 1 mg wet tissue (100 μg protein) was reduced with 10mM dithiothreitol at 56 °C for 30 min and alkylated with 10 mM iodoacetamide at room temperature in the dark for additional 30 min in ultrafiltration centrifuge tube (Millipore, USA). Then samples were followed the FASP method digestion with trypsin (MS Grade, Pierce™, Thermo Fisher Scientific, USA). The combined filtrates were desalted on a high pH reversed-phase peptide fractionation column (Pierce™, Thermo Fisher Scientific, USA). Then the peptides were collected followed by vacuum drying. The chemicals used for protein extraction and digestion were of MS Grade.

### LC-MS/MS for Proteomics Analysis

The collected peptide samples were analyzed by an Easy nLC Orbitrap Fusion Lumos platform (Thermo Fisher Scientific, USA). About 100 μg peptides were loading for the proteomic label free quantification. The dried peptide samples were re-dissolved in 0.1% formic acid and loaded onto a C18 column (100 μm× 20 mm, 3 μm, Thermo Fisher Scientific, USA) then separated on a C18 analytical column (150 μm × 120 mm, 1.9 μm, Thermo Fisher Scientific, USA) with a gradient of 7-40% mobile phase B (80% acetonitrile and 0.08% formic acid) at a flow rate of 600 nl/min for 70 min. The master scan range was 300 to 1,400 at a resolution of 120,000 at m/z 200. The automatic gain control (AGC) target was set at 500,000. The maximum injection time was set at 50 ms. Included charge state in filter charge state was 2 to 6. Polarity was positive. Data-dependent mode was used for MS2 scan. Isolation window was 1.6. First mass was from 120. The precursor was subjected to fragmentation via higher energy C trap dissociation (HCD) with 30% normalized collision energy. The AGC was set at 5,000 and the maximum injection time at 35 ms. The Xcalibur software was used for acquiring data (Thermo Fisher Scientific, USA).

### OA Proteome Quality Assessment

Distribution of relative standard deviation (RSD) in each group was used to describe the quantification stability in OA study during the LC-MS/MS analysis. The mean value of RSD for all the quantified proteins were 0.428, 0.496, 0.461, 0.433, and 0.479 in control, OA model, GA, ESDL, and ESDH group, respectively. The closed RSD results indicated that the data of quantitative protein among different samples in the same group had little variation, while using the label free quantitative method to calculate the relative quantitative value according to the iBAQ value. The SD range of RSD in each group was between 0.415 and 0.474, and the variation was not significant. The results showed that the total variation degree of quantitative results of different proteins in each group was similar. Therefore, the work flow had little variation and could be used for pharmacological effect analysis and evaluation.

### Mass Spectrometry Data Analysis and Bioinformatics Analysis

All raw files were submitted to j MaxQuant software (version 1.6.3.4) and compared against the UniProt rat protein database (version 20180903, 8,033 sequences). The target-decoy based strategy was used to achieve a peptide and protein false discovery rate (FDR) ≤ 1%. The search parameters were as described previously (22). Decoys for the database search were generated with the revert function. The following options were used to identify the proteins: Peptide mass tolerance 15 ppm, MS/MS tolerance 0.02 Da, digestion enzyme trypsin, missed cleavage 2, fixed modification: Carbamidomethyl (C), variable modification: oxidation (M). The selection criteria of differentially expressed proteins was unique peptide ≥ 2, *p* ≤ 0.05, and identified in at least 33.3% of total samples (≥8). Gene Ontology annotation was searched against DAVID (23). Pathway and protein-protein interaction were performed using STRING database (24).

### Osteoarticular Cartilage Histopathology

Following osteoarticular cartilage collection, tissues were fixed with 10% neutral-buffered formalin. To evaluate inflammatory cell infiltration, each group was stained with hematoxylineosin (H&E) and safranine O-fixation green staining. The instrument was 80i biological microscope (Nikon, Janpan), VIP-6 totally closed tissue dehydrator (Cherry Blossom, Japan), Rm2235 microtome (Leica, German), Dyeing and sealing machine (Cherry Blossom, Japan).

### Pressure Pain Threshold (PPT)

The PPT was measured by a pressure algometer. After intragastric administration, the right knee joint of the rat was put on the platform of pressure algometer. The tenderness instrument stopped automatically when the rat struggles, then the test stopped, and the pain threshold was recorded.

### Electronic Von Frey Test

Von Frey fillings is composed of 20 nylon fibers, which can provide 0.008g-300g ciliary mechanical stimulation. It can be used for quantitative pain test of a variety of animals (20). In our study it was used to estimate the mechanical pain threshold of injured rats’ knee joint by Von Frey hair test. The rats were placed on a transparent metal frame with a glass box cover and with mesh holes on the bottom. After a period of adaption, when the rats were quiet, a series of ascending forces of electronic von Frey single filament were used to stimulate the outer side of the right rear paw of the rats. The ascending force gradually increased from light to heavy, until a sharp retraction of the right paw. Then the stimulating figures were recorded as reflex pain threshold.

### Measurement of knee joint curvature

In each group, the rats were anesthetized routinely, using the cotton swab to locate parallel to the edge of the working table. The femoral end of the anesthesia rats was fixing to align with the cotton swab. The lower limbs were pulling with same force for abduction and adduction and stopping in case of slight resistance.

### Enzyme-Linked Immunosorbent Assay (ELISA)

Blood samples were collected in test tubes and centrifuged for 15 minutes to collect serum at 3500 rpm. Serum levels of TNF-α, IL-1β, and IL-6 were measured using a commercially available ELISA kit (R & D, Minneapolis, USA), following the manufacturer’s instructions.

### Quantitative Real-time PCR RT-qPCR) Analysis

Osteoarticular cartilage tissues were used for real-time RT-qPCR. Total RNA was extracted with RNeasy^®^ Lipid Tissue Mini Kit (QIAGEN, California, USA). RNA quality and quantity were determined using a spectrophotometer (Thermo Nano DropTM 2000c, USA) and denaturing agarose gel electrophoresis. The ApoE, C3, C16, p38, Akt, and GAPDH genes were amplified using the forward and reverse primers listed in supplementary table 14. The RNA (2μg) was used for cDNA synthesis using reverse transcription master Mix (QIAGEN, California, USA). RT-qPCR was performed in SYBR Green PCR Master Mix (QIAGEN, California, USA) on CFX-96 system. To normalize the sample variance, the GAPDH gene was used as an endogenous control. The experiment was repeated three times.

### Western blot analysis

Approximately 50 mg of cartilage tissue from each rat was used to extract protein using RIPA buffer containing PMSF and Phosphatase inhibitor. Protein concentration was determined by BCA assay, and total proteins were separated by 8-10% SDS-PAGE. 10 μg protein was loaded per lane of a 1mm thick mini polyacrylamide SDS-gel. The separated proteins were then transferred to PVDF membranes (0.45μm, Millipore, USA), and block with 5% milk for 1h. After blocking, the membranes were washed three times with TBST, and incubated with primary antibodies including ApoE (ab183597), p38 (ab31828), p-p38 (phosphoY182, ab47363), AKT1+AKT2+AKT 3(ab179463), anti-AKT1 (phosphoS473, ab81283), C6 (proteintech17239-1AP), complement C3 (ab200999) (1:1000 diluted), *β*-Actin (proteintech66009-1-lg) and GAPDH (1:5000 diluted) overnight at 4 °C, and incubated with the secondary antibody (1:10,000 diluted) for 1 h at room temperature to detect antibody binding. Protein bands were quantified using a ChemiDoc™ MP Imaging system (Bio-Rad Co., USA), and analyzed using the Image Lab software (Bio-Rad Co., USA). Finally, values were expressed as band intensity normalized to GAPDH or *β*-actin.

### Statistical analysis

The statistical significance of the differences between groups was determined by ANOVA. All the data was shown by mean ± SD and *p*≤0.05 was considered significant.

## RESULTS

### The ESD Component Identification by LC-MS

The LC-MS for determination of the components of ESD was shown in figure 1B. We found twelve compounds in ESD through mass spectrometry peaks (Fig. 1C). The identified compounds included Scutellarein 5,6,7,4’-tetramethyl ether Tetramethylscutellarein ([M+H] m/z 343.1729, 4.7 min), Icariin ([M+H] m/z 677.2432, 5.2 min), Epimedin C ([M+H] m/z 823.3015, 5.2 min), 8-Prenylquercetin 4’-methyl ether 3-rhamnoside ([M+H] m/z 531.1864,8.2 min), Ferulic acid ([M+H] m/z 194.1175, 9.2 min), 1,2-Dihydrotanshinone ([M+H] m/z 279.1014, 9.8 min), Diosgenin acetate ([M+H] m/z 457.4473, 9.8 min), Cryptotanshinone ([M+H] m/z 297.1481, 10.4 min), Tanshinone I ([M+H] m/z 277.0857, 10.4 min), Tanshinone IIA ([M+H] m/z 295.1325, 11.4 min), Delta3,5-Deoxytigogenin ([M+H] m/z 413.3358, 11.4 min), Salvirecognone ([M+H] m/z 285.2893, 14.4 min) (supplementary table 1).

**Figure 1.**
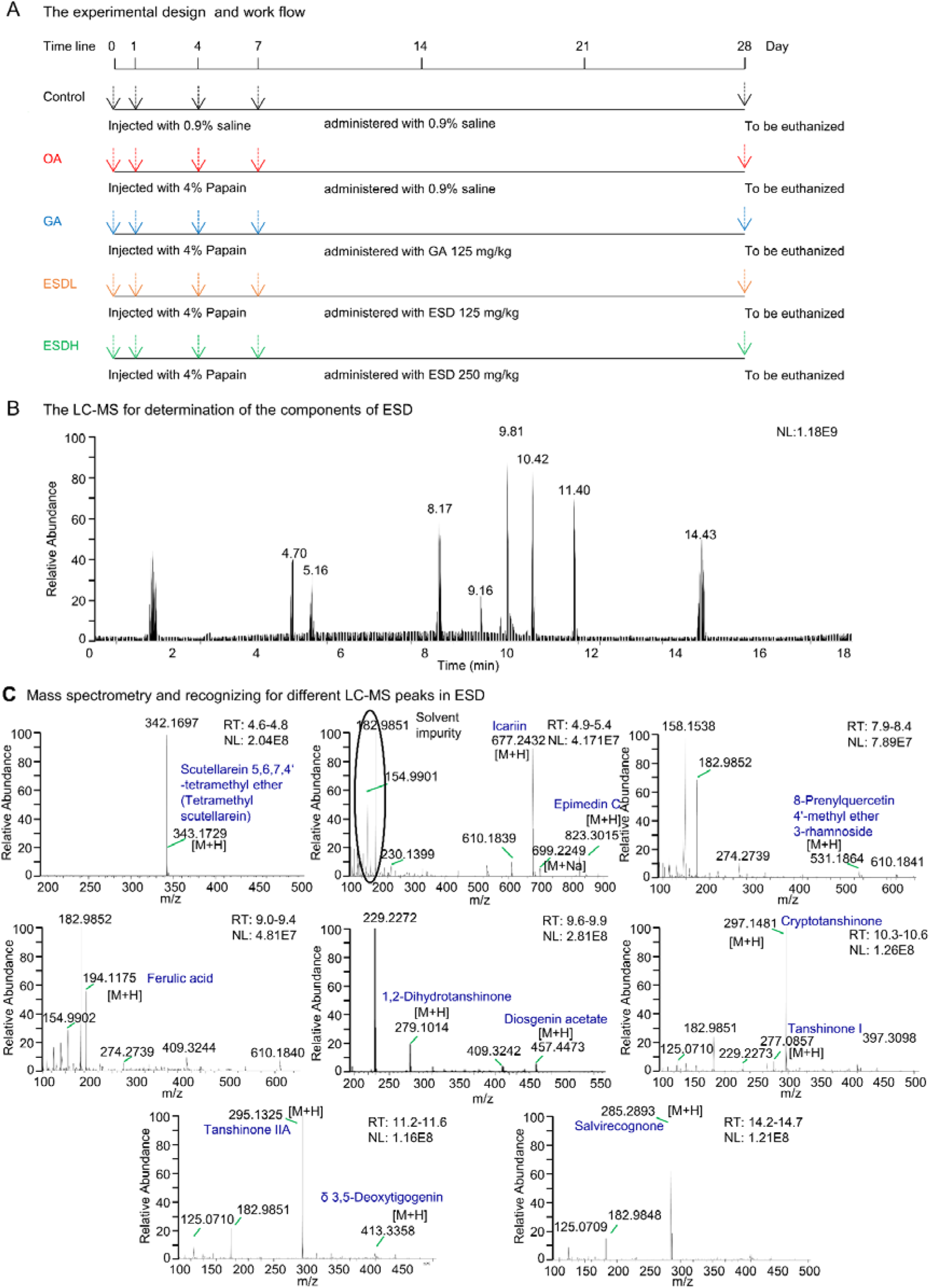
A. Workflow of papain-induced rat model and ESD administration. B. The LC-MS for determination of the components of ESD. C. Mass spectrometry and recognizing for different LC-MS peaks at different RT in ESD (RT: retention time; NL: target level).

### General Situation and Behavioral Observation

There was no death and no significant difference in body weight between groups during the entire process of the study. The rats in normal control group were in good mental state during entire evaluation process. No abnormal activity was found. The diet and drinking water in control group were normal. In the later stage of the evaluation, the rats in OA group showed decreased autonomic activity and increased lying down and laziness compared with control group. There were redness and swelling phenomenon in the right limb knee joint of model group. As the administration time prolonged, the autonomic activity of rats in both ESDL and ESDH groups were slightly better than that of the model group.

### General Features of the OA Proteome

Seeking to gain insight into OA, we used mass spectrometry to identify proteins differently expressed in the articular cartilage of five groups. The workflow was showed in figure 2A. The peptide spectrometry match (PSM), identified peptides, and proteins were 286325, 13199 and 1543 respectively (supplementary table 2). The quantified proteins number was 1537 (unique peptides (UP) ≥1), 1444 (UP≥2), and 1408 (UP≥2, and identified in at least 8 samples) (Fig. 2B, supplementary table 3,4). The change protein numbers were showed in figure 2C. The overlap of down-regulated proteins (OA *vs*. control) and up-regulated proteins (GA *vs*. OA, ESDL *vs*. OA, and ESDH *vs*. OA) were 97, 124, and 78, respectively (Fig. S1A, supplementary table 5-12). Distribution of relative standard deviation (RSD) in three independent pooling samples of all groups was shown in figure 2D. The RSD value of different groups was close to each other, range from 0.4 to 0.5, which indicated the good stability of the whole experimental flow. The changed proteins in OA *vs*. control were shown by volcano map (Fig. 2E). Using a stricter cut off criterion, the number of down-regulated proteins was more than the up-regulated in OA group compared with control (66 *v.s*. 12).

**Figure 2.**
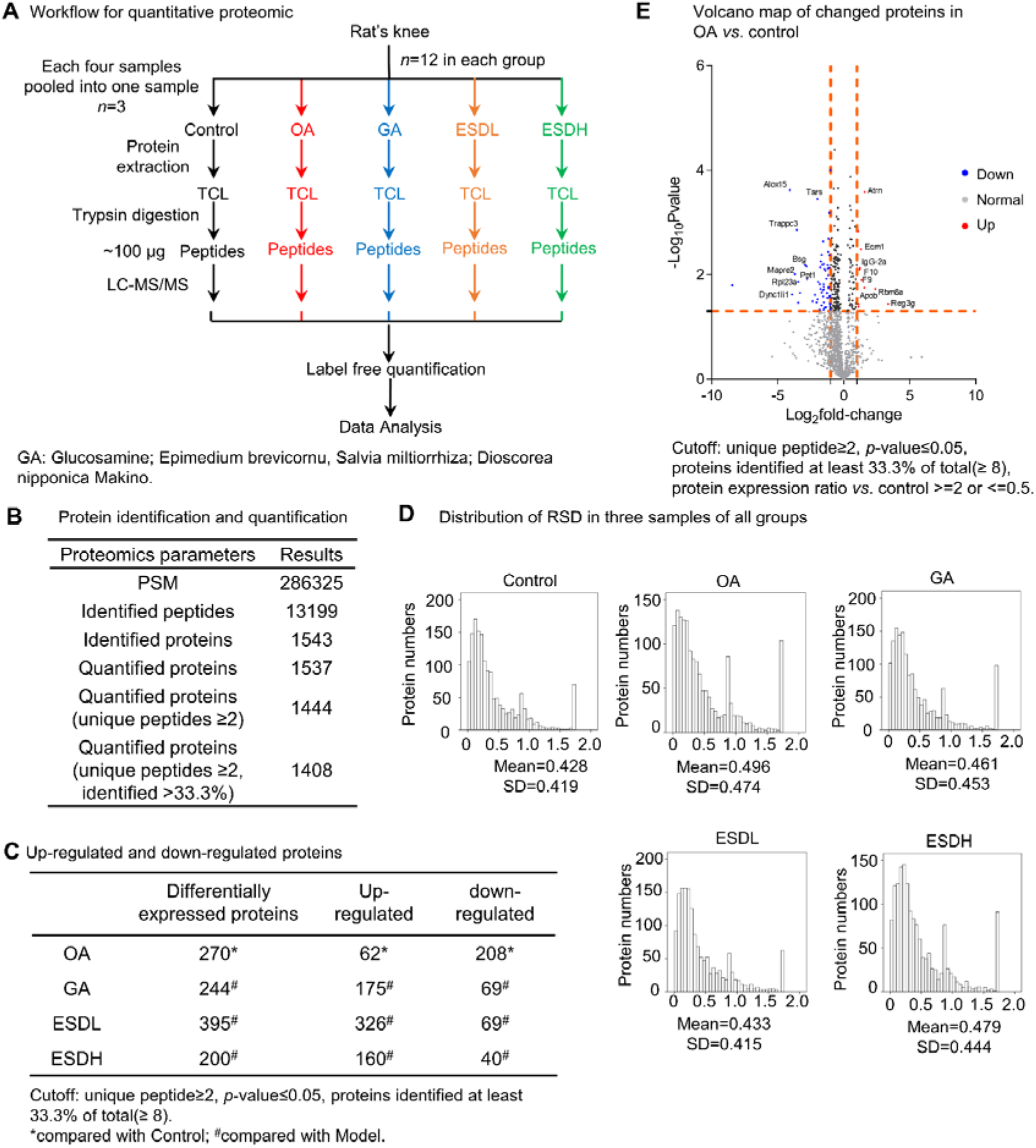
A. Workflow for quantitative proteomic study. B. Protein identification and quantification. C. Up-regulated and down-regulated proteins for each group. Cutoff: unique peptide≥2, *p*-value≤0.05, and proteins identified at least 33.3% of total samples (≥ 8). D. Distribution of RSD in three pooling samples of all groups. E. Volcano map of changed proteins in OA vs. control. Cutoff: unique peptide≥2, *p*-value≤0.05, proteins identified at least 33.3% of total (≥ 8), protein expression ratio vs. control ≽2 or ≼0.5.

### Humoral Immune Response and Complement Activation Up-Regulated in OA and ESD Restored the Expression of Changed Proteins

Through PCA analysis we found the points of the control, GA, ESDL, and ESDH groups were distributed nearby, and the OA group was far from the other groups (Fig. 3A). The PCA analysis results explained more than half of the variation (about 63.6%, Dim1 add Dim2) among the total fifty samples. Ratio distribution of OA model up-regulated proteins in different five groups were compared and the curves shifted to the left by drug treatments (Fig. 3B). Sting analysis showed the up-regulated proteins in OA model group were involved in the biological process of immune response, activation of humoral immune response, complement activation, leukocyte mediated immunity, acute inflammatory, regulation of endocytosis, regulation of proteolysis (Fig. 3C, D, supplementary table 5). The complement activation proteins included Complement C3, Complement C5, Complement C4A, SERPING1, Clusterin, and Ficolin-2. From these up-regulated proteins in OA model and restored in ESD group we find the in vivo cellular targets for ESD were regulation of immune response and cellular metabolic process. The changed of complement C3 and C6 in each group were shown in figure 3E. The complement level of C3 and C6 were significantly up-regulated in OA model and restored after GA and ESD administration. Plasma lipoprotein particle remodeling proteins were also involved in OA injury (Fig. 3F). Apolipoprotein B (ApoB) and Apolipoprotein C-I (Apo C-I) were significantly up-regulated in OA group and restored after ESD administration. Oxidative stress related proteins such as Cu-Zn superoxide dismutase and glutathione peroxidase 3 were up-regulated in OA model group and restored in ESD administration groups (Fig. 3G).

**Figure 3.**
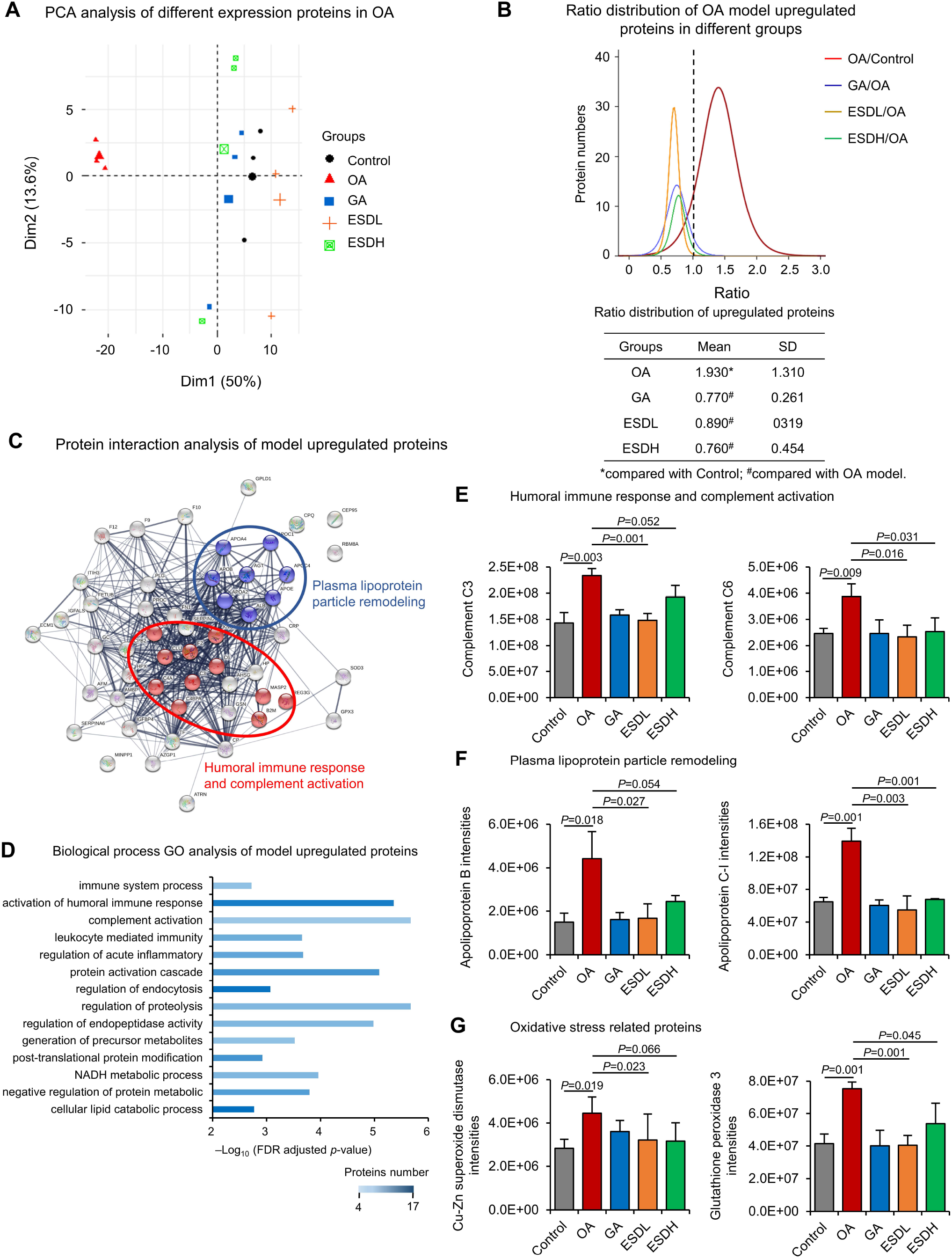
A. Score plot from PCA based on all detected proteins. Apart from group “OA” (red), all replicates cluster together and the different groups (i.e., control, GA, ESDL, and ESDH) cluster away from each other, visualizing the differences between the groups. B. Ratio distribution of arthritis model up-regulated proteins in different groups. C. Protein interaction analysis of model up-regulated proteins through string database analysis. D. Biological function GO analysis of model up-regulated proteins. E. Level of humoral immune response and complement activation related proteins in each group. F. Level of Plasma lipoprotein particle remodeling related proteins in each group. G. Level of oxidative stress related proteins in each group. Note: Data are expressed as mean ± SD (E-G, pooling samples, *n*=3).

### Leukocyte Mediated Immunity and Mapk Signaling Pathway Negative Regulation Proteins were Down-Regulated in OA Model Group

The mean value of Log2ratio (OA/Control), Log2ratio (GA/OA), Log2ratio (ESDL/OA), and Log2ratio (ESDH/OA) were −0.990, 0.790, 1.000, and 0.840 respectively (Fig. 4A). It suggested that ESD increased the down-regulated proteins in OA model. The ESDH group has the highest Log2ratio value according with the most remarkable protective effect. The down-regulated proteins in OA model group were involved in the biological process of metabolic proteins, leukocyte/neutrophil mediated immunity, leukocyte activation, and MAPK signaling pathway negative regulation proteins (Fig. 4B, supplementary table 6). The metabolic process proteins were involved cofactor metabolic process, carboxylic acid metabolic process, carboxylic acid metabolic process, coenzyme metabolic process, purine nucleotide metabolic process. The leukocyte mediated immunity activation was down-regulated, which was opposite to the trend of humoral immune response (Fig. 4C, S1B). Muscle contraction and muscle filament sliding proteins were decreased significantly which correspondence with the motor function decline caused by OA.

**Figure 4.**
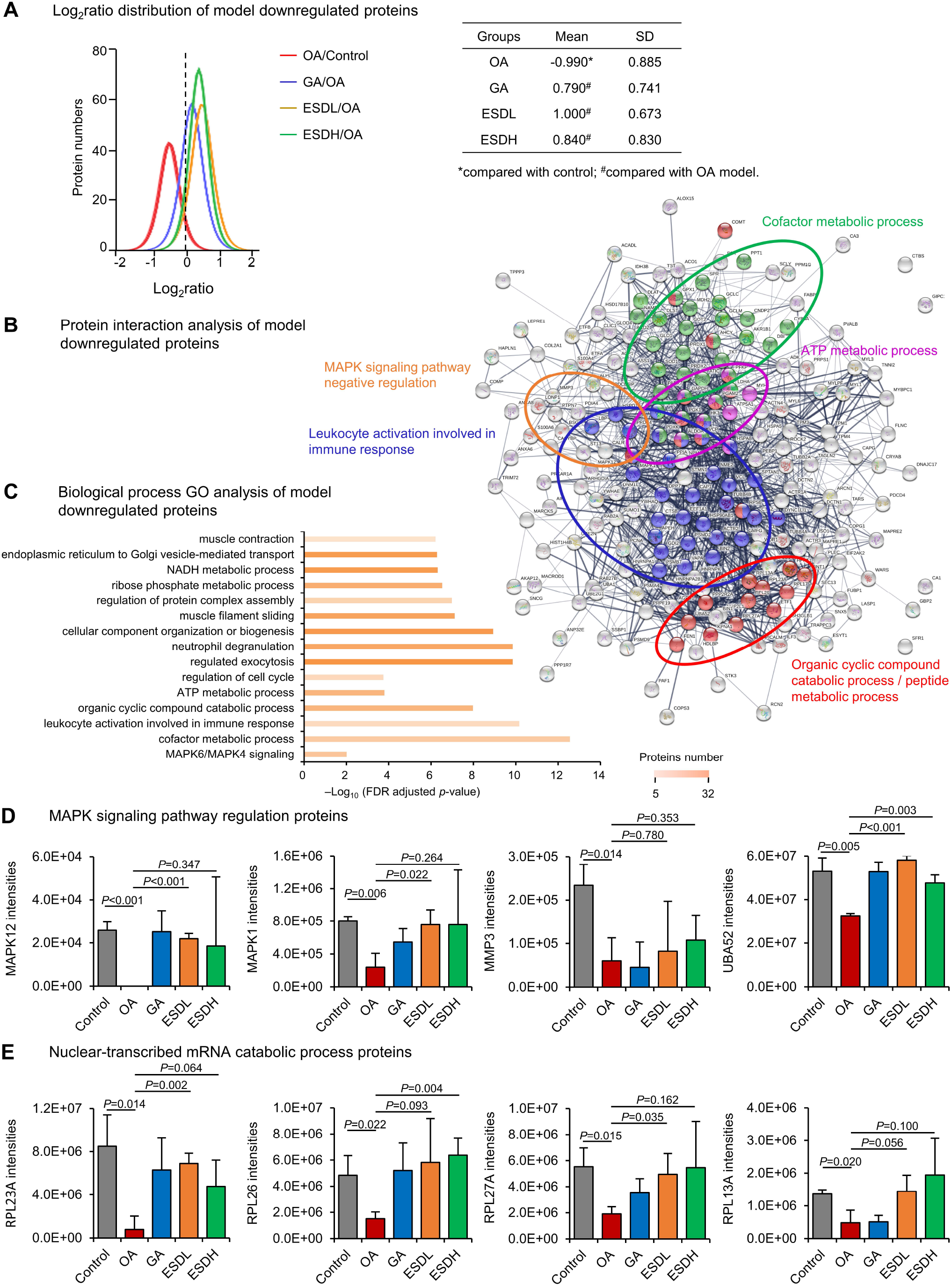
A. Log2ratio distribution of OA model down-regulated proteins. B. Protein interaction analysis of model down-regulated proteins. C. Biological process GO analysis of model down-regulated proteins. D. MAPK signaling pathway regulation proteins’ intensities. E. Nuclear-transcribed mRNA catabolic process proteins. Note: Data are expressed as mean ± SD (D-E, pooling samples, *n*=3).

The MAPK signaling related proteins were significantly down-regulated in OA model and reversed after ESDH administration. These proteins included Mitogen-activated protein kinase 12 (MAPK12), Tyrosine-protein phosphatase non-receptor type 7 (PTPN7), Basigin (BSG), cAMP-dependent protein kinase type I-alpha regulatory subunit (PRKAR1A), Stromelysin-1 (MMP3), Mitogen-activated protein kinase 1 (MAPK1), and (UBA52) (Fig. 4D). Among them, MAPK1 and UBA52 were involved in negative regulation of MAPK pathway.

### Ribosomal Proteins were Down-Regulated in OA Model and Restored by ESDH

In our results we found ribosomal proteins down-regulated such as the arthritis-related autoantigen 60S ribosomal protein L23A (RPL23A) (25), ribosomal protein L26 (RPL6), ribosomal protein L27A (RPL27A), and ribosomal protein L13A (RPL23A), ribosomal protein L17 (Rpl17), Ubiquitin-60S ribosomal protein L40 (Uba52), and 40S ribosomal protein S6 (Rps6) were significantly decreased in model group, and partially restored after ESDH administration (Fig. 4E).

### Heatmap analysis and HCL clustering

In heatmap HCL cluster distance metric selection part, Pearson correlation was chosen as current metric, and average linkage clustering was used as linkage method selection. We got eight clusters by HCL clustering calculation (Fig. 5A, supplementary table 13). Cluster 1 included proteins involved in regulation of plasma lipoprotein particle levels were APOC1, APOA4, APOC4, APOB, APOA2, APOE, ALB, AGT, CRPP, and C3. Proteins involved in regulation of proteolysis were C6, C9, FETUB, F12, ITIH3, CLU, AHSG, AMBP, SERPINA6, C3, APOE, FN1, GSN, and AGT. Cluster 3 included leukocyte activation and hydrolase activity related proteins. Leukocyte activation proteins included DPP7, MAPK1, PDXK, BIN2, HSPD1, DYNC1LI1, RPL13A, PSMD2, PSMB7, and APEH. Hydrolase activity related proteins included MMP3, DPP7, MYL3, PPM1G, FEN1, DYNC1LI1, ACP1, APEH, PSMB7, PSMA4, and PSMD2. Cluster 4 included complement cascades proteins and proteolysis regulation proteins. Cluster 5 included acetylation and phosphoprotein proteins, cellular responses to stress, and proteins of cellular response to cytokine stimulus. Cluster 6 included acetylation and phosphorylation proteins and metabolism of proteins. Cluster 7 included muscle system process and muscle contraction proteins, small molecule metabolic process related proteins, and leukocyte mediated immunity proteins. Cluster 8 included small molecule (purine nucleotide) biosynthetic process proteins, anatomical structure development proteins, response to stress, and metabolic pathways related proteins.

**Figure 5.**
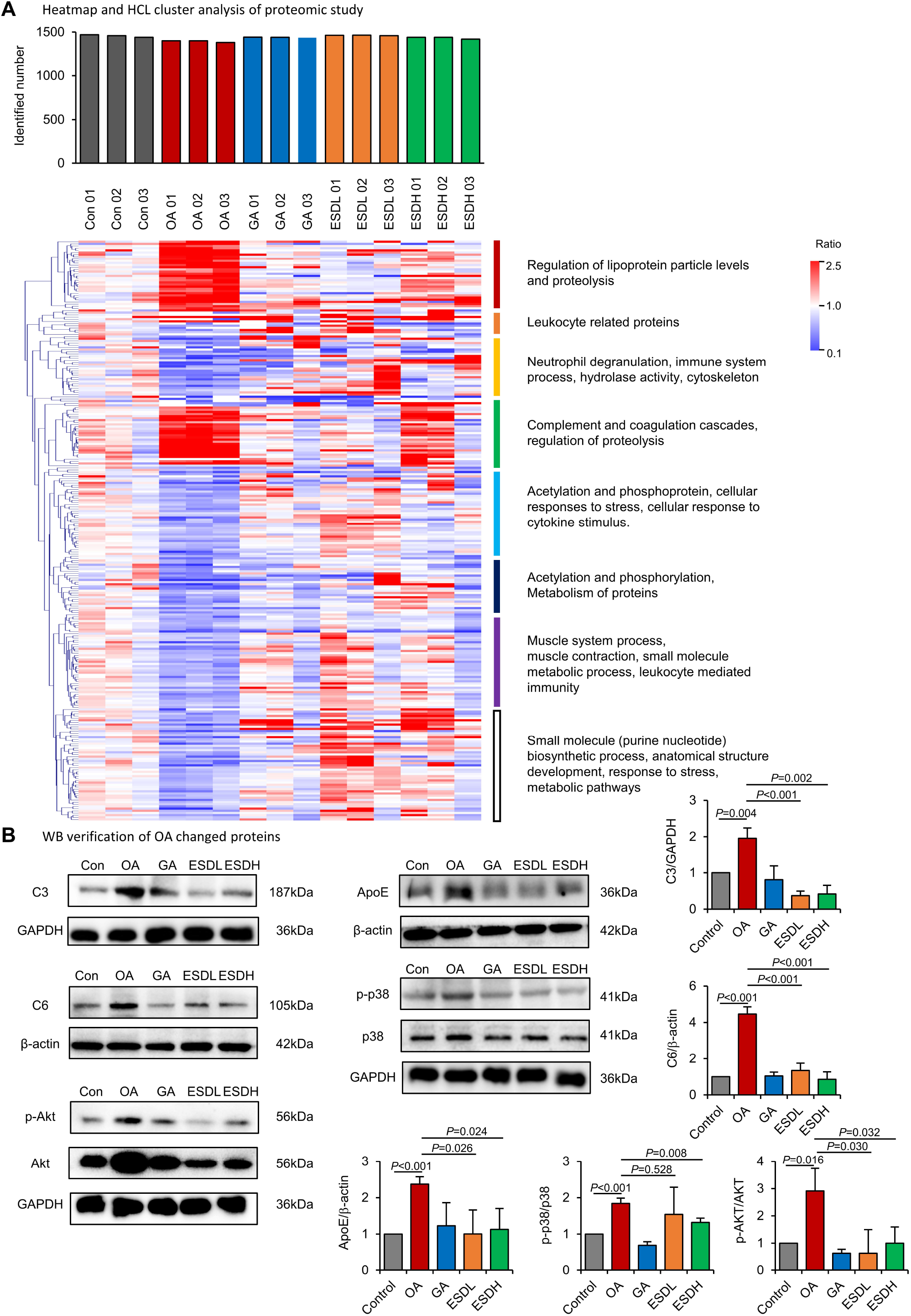
A. Heatmap analysis and HCL clustering of OA changed protein in all samples. There were eight cluster with different function and expression levels in five groups. B. PCR and WB verified the change proteins. Note: Data are expressed as mean ± SD (B, pooling samples, *n*=3).

### PCR and WB Verified the Change Proteins of Proteomics’ Results

The results of PCR and WB were consistent with those of proteomics (Fig. 5B, S1D, S2). We found C3 and C6 in OA group were significantly up-regulated in both gene and protein levels. The levels of C3 and C6 in GA and ESD groups decreased to close to normal level. ApoE gene and protein level in OA group increased significantly. Apolipoprotein ApoE level in GA and ESD group decreased to near normal level. Akt pathway related proteins Akt and p38 were up-regulated in OA group in both gene and protein levels. Akt and p-p38 levels in GA and ESD groups were decreased to close to normal levels.

### ESD Improved Morphological Changes of Articular Cartilage in Rat Lower Femur

HE staining and safranin O-fast green staining was used to study the morphological changes of articular cartilage. In control group, articular cartilage was normally divided into surface layer, transitional layer, radiation layer and calcification layer. Chondrocytes were arranged in order and the tide line was complete. In OA group, the surface of articular cartilage was irregular, and lot of cracks were seen in the field of microscope (Fig. 6A). The local cracks were deep to the calcified layer. The surface layer of cartilage was peeled and the peeling range was large. Free cartilage tissue was seen in articular cavity. Similar with OA group, the surface of articular cartilage in GA group was irregular, with many cracks. The articular cartilage extended to the articular cavity in form of villi. The local cracks were deep to the calcified layer, and the surface layer of cartilage peeled off. The “cluster like” chondrocyte clusters appeared. In ESDL group, the articular cartilage surface of some animals was irregular, the cartilage layer became thin. The cracks were common, but most of them were located in the superficial layer. In ESDH group, the articular cartilage surface of some animals was irregular, with many cracks, mostly in the shallow layer. Some cracks were extending to the joint cavity in form of villus. The local cracks were deep to the calcified layer. The surface layer of cartilage was peeling off. The peeling range was large, with free cartilage tissue in the joint cavity. There were different degrees of injury in OA and each group but control. ESD has a certain protective effect, and its low dose group has the most obvious protective effects.

**Figure 6.**
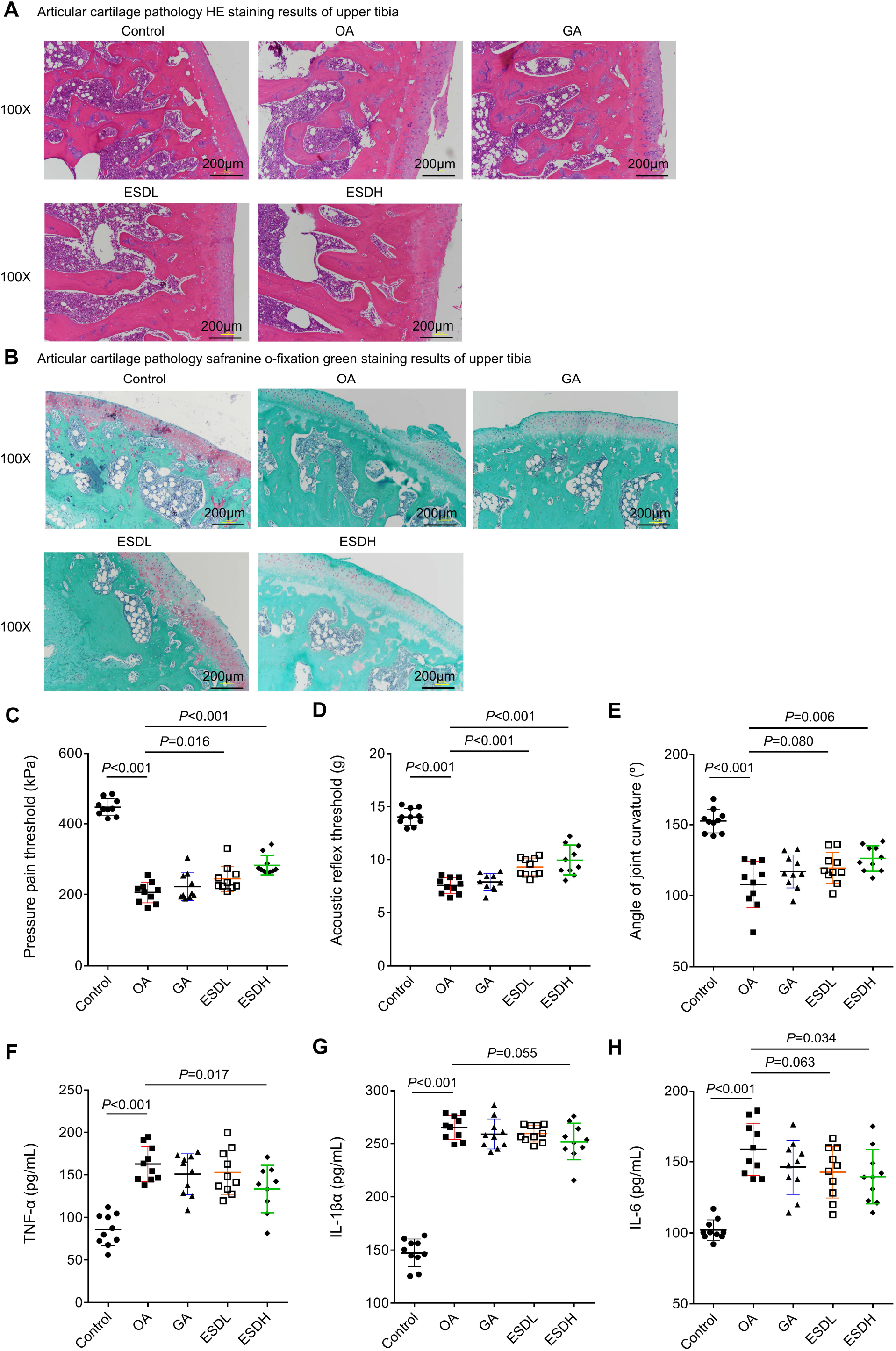
A. H&E staining results of articular cartilage of lower femur (100×). B. Safranin O-fast green staining of articular cartilage of lower femur (100×). C. The pressure pain threshold for different groups after OA modeling and GA or ESD administration. D. The reflex pain threshold for different groups in Von Frey hair test. E. Measurement of knee joint curvature for different groups. F to H. The level of serum TNF-α, IL-1β, and IL-6 concentration in different groups. Note: Data are expressed as mean ± SD (B, pooling samples, *n*=10).

Cartilage consists of cartilage tissue and perichondrium. Cartilage can be divided into hyaline cartilage, elastic cartilage and fibrocartilage according to the cellulose content in the matrix. The structure of cartilage, subchondral bone, osteogenic tissue and tidal line of knee joint in rats can be displayed by safranine O-fast green staining. Among them, the cartilage is red or orange red, and the osteogenesis is green, which is in sharp contrast with the red cartilage, so as to distinguish the cartilage and bone tissue. Safranine O-fast green is a kind of cationic dye which combines with polyanion. The showing of cartilage tissue is based on the combination of cationic dye and anionic group (chondroitin sulfate or keratin sulfate) in polysaccharide. When cartilage was damaged, glycoprotein in cartilage was released, resulting in uneven distribution of matrix components. So the Safranine O-fast green staining was light or stainless. In control group, the staining of articular cartilage matrix was basically normal, uniform red, and the staining around chondrocytes was deepened, while the staining far away from chondrocytes was lighter (Fig. 2B). In OA group, the articular cartilage matrix was stained shallowly, unevenly and even could not be stained. The cartilage matrix of GA group was stained shallowly or unstained. The cartilage matrix of ESDL group was slightly stained. In ESDH group, the staining of articular cartilage matrix was shallow and uneven.

### ESD Down-Regulated OA Pressure Pain Threshold (PPT)

Pain threshold is the minimum intensity that a stimulus as being painful, and is assessed by a series of ascending and descending stimulation intensities (26). We found the right posterior knee tenderness threshold in the OA group was significantly lower than that in the control group. This indicated that while closing to the test end point, OA lead to raising sensitivity to pressure pain. The mean value of PPT (kPa) for control, OA, GA, ESDL, and ESDH were 447.8±22.7, 205.8±27.4, 222.4±36.8, 244.5±34.3, and 283.1±26.7 (mean±SD) respectively. The right posterior knee PPT of the model group was significantly lower than that of the normal group (Fig. 6C). ESDL and ESDH significantly decreased the sensitivity of rats’ knee joint and restored PPT. PPT of GA group was slightly increased than that of model, but with no significant difference.

### ESD Restored OA Reflex Pain Threshold in Electronic Von Frey Test

The mean value of prickling reflex threshold (g) for control, OA, GA, ESDL, and ESDH were 14.03±0.76, 7.53±0.70, 7.86±0.75, 9.30±0.80, and 9.94±1.34 (mean±SD) respectively (Fig. 6D). The right posterior knee reflex threshold of the model group was significantly lower than that of the normal group. ESD significantly restored the reflex pain threshold in both lower and higher dosage. The reflex pain threshold of GA group was slightly higher than that of model, but with no significant difference.

### Measurement of Knee Joint Curvature

We measured the angle between the two action lines and the cotton swab as the knee joint curvature of the rats. The mean value of knee curvature (°) for control, OA, GA, ESDL, and ESDH were 152.7±7.9, 107.9±15.6, 117.1±11.2, 119.5±10.5, and 126.3±8.5 (mean±SD), respectively (Fig. 6E). Right posterior knee curvature in OA group was significantly decreased compared with normal control group. ESDH significantly restored knee curvature compared with OA.

### Determination of Serum Inflammatory Factors

TNF-α, IL-1β, IL-6 are mainly involved in immune regulation, infection and inflammatory response in the body (27). They are mainly secreted by Th1 type cells, and are also the marker cytokines of type I helper T cells (Th1 cells). In our results, the concentrations of TNF-α (Fig. 6F), IL-1β (Fig. 6G), IL-6 (Fig. 6H) was significantly higher in serum from individuals with OA group and each administration group than in serum from normal individuals. Thus, inflammatory activation occurs in blood in the course of OA and persist. ESD significantly decreased concentrations of TNF-α, IL-1β and IL-6 in both lower and higher dosage compared with control group. These results of serum inflammatory factors were consistent with proteomics analysis.

## Discussion

TCMs have long been used to treat OA-related diseases. Identification the key components and underlying molecular pharmacological mechanisms remains a challenge for multiple herbal constituents TCM study. Proteomic and bioinformatics have been developed as novel methods to indicate the pharmacological and biological mechanism of TCM. This study we investigated the protective effects of EDS extracts in mice OA model. We also presented a proteomic landscape of OA and EDS treatments, suggesting that ESD partially restored OA-induced protein change in rats’ model.

Twelve main components were identified in the positive and negative-ion mode through mass spectrometry. These compounds allow us to gain insight of main ingredients of ESD. The identified compound scutellarein 5,6,7,4’-tetramethyl ether was reported to suppress lipopolysaccharide (LPS)-induced cyclooxygenase-2 (COX-2) and inducible nitric oxide synthase (iNOS) via inhibition of the NF-κB pathway (28). Icariin was reported to alleviate OA by inhibiting nod-like receptor protein-3 (NLRP3)-mediated pyroptosis (29). It also inhibited cartilage degradation by decreasing metalloproteinase-13 (MMP-13) expression in IL-1β-induced chondrocytes and in an experimental OA model (30). Icariin and Epimedin enhanced alkaline phosphatase (ALP) activity, mineralization, and osteoblasts proliferation in LPS-induced osteoblasts. *In vitro* evaluation showed that epimedin C and icariin significantly promoted ALP activity in osteoblast-like cells. The bioactivity of epimedin C and icariin may contribute greatly to prevention of ovariectomised-induced rat model bone loss (31). Ferulic acid was reported to alleviate the symptoms of OA (32). The ESD’s protection against OA might come from the anti-inflame effects of the identified compounds.

The changed proteins were located to the same signaling pathway or played the similar biological function according to the proteomic results, which indicated the reliability of the proteomic results. The number of down-regulated proteins was far more than the up-regulated proteins in OA group compared with control group. The number might indicate the function damage of OA rats’ knee in model group. The PCA results indicated that the GA, ESDL, and ESDH groups were all showed some potential protective effects for OA. Most of the model group down-regulated proteins was up-regulated in ESDL group, which may explain the partial improvement of movement function of OA knees. The restored changed proteins might contribute to the restored movement function of knee joint OA. Partial of the model group down-regulated proteins was up-regulated in ESDH group. The restored proteins included complement cascades, leukocyte mediated immunity, regulation of plasma lipoprotein particle levels, and muscle contraction or muscle filament sliding. The activation of complement may be the cause of proteolysis level in OA group, which resulted in the function damage of knee movement.

From these up-regulated proteins in OA group and restored in ESD group we found the in *vivo* cellular targets for ESD were regulation of immune response and cellular metabolic process. Inflammatory complement system plays a central role in the pathogenesis of OA (33). The up-regulated complement level could increase the proteolysis of cartilages, which partially explained the movement function damage of knee joint. The upregulation of complements in OA was consisted with previous report (33, 34). We found that expression and activation of complements was abnormally high in OA group, initiating proteolysis of knee cartilage. Expression of inflammatory and complement proteins was lower from ESD treatment groups, indicating that ESD has certain therapeutic effects in rats. The similar trend of C3, C4A, and C5 in our proteomic results once again demonstrated the crucial role of complement to the development of arthritis in mouse models of OA. ESD’s protective effect partially showed by the restoration of levels of complements. As inflammatory complement system played a central role in the pathogenesis of OA, ESD down-regulated the activation of complements in humoral immune system thus to fulfill the protection against OA damage. The activation of complement may be the cause of proteolysis level in OA group, which resulted in the function damage of knee movement.

A recent porcine knee synovial fluid proteome study found apolipoprotein A4 (ApoA4), Apolipoprotein C3 (ApoC-III), and Apolipoprotein E (ApoE) were detected as abundant proteins in the synovial fluid in the porcine posttraumatic OA model. In our study, we found ApoA-II, ApoB, ApoC-I, ApoC-IV, ApoE, and ApoA-IV were significantly up-regulated in OA group and restored after ESD administration (supplementary table 5). Among them, ApoB was reported as a key component of atherogenic lipids. Our result was consisted with previous report that ApoB aggravated inflammation in rheumatoid arthritis (35). ApoB aggravated arthritis by potentiating the inflammatory response via its interaction with enolase-1 expressed on the surface of immune cells. ESD protected articular cartilage anddecrease the injury of OA by downregulating the expression of ApoB (Fig. 3F). Another apolipoprotein (AopA-I) was also found to be decreased after Ultrasound (US) therapy in a rabbit knee OA model from a synovial fluid proteome study, which might be related with the OA symptoms improvement (36). Apo A-I was identified in amyloid deposits in knee joints in patients with knee OA (37). The moderation effect on level of apolipoprotein may partially explain the protection of ESD.

The leukocyte mediated immunity activation was down-regulated, which was opposite to the trend of humoral immune response (Fig. 4C). The down-regulated leukocyte activation included more than 30 proteins, which was far more than that of up-regulated humoral immune response proteins (Fig. S1B). Many important biological process proteins were down-regulated in OA group, e.g. cytoskeleton organization, the biological process of metabolic proteins, even including cellular immune related proteins. Leukocyte mediated immunity and MAPK signaling pathway negative regulation proteins were down-regulated in OA group. These results indicated that the leukocyte immune response might be depressed while humoral component response was activated. Muscle contraction and muscle filament sliding proteins were decreased significantly which correspondence with the motor function decline caused by OA. These results partially explained the damage of OA and protective effect mechanism of ESD. The results indicated that the damage of OA might be mainly form the complement in humoral immunity other than cellular immunity.

Previous studies also reported 60S ribosomal protein L10-like and 60S ribosomal protein L18 were found down-regulated in rheumatoid arthritis (38). Actually, we found lot of ribosomal proteins down-regulated. Among them, Rpl23a was described as one immune dominant protein for the local T cell response in reactive arthritis (39). These results showed the ribosome function injury and protein biosynthesis dysfunction during arthritis process. ESD has protective effects for the ribosomal protein in both low and high dose.

## CONCLUSIONS

In conclusion, our study provided proteomics evidence for ESD alleviated OA through the modulation of protein expression. *In vivo* studies showed ESD relieved symptoms from OA cartilage degradation. The protein profiling protective mechanism for ESD was through attenuating inflammation pathways and modulating humoral and cellular immune response. The present work provided label-free quantitative proteomics solution for verifying the protective effects of potential multi-components OA medicine.

## Acknowledgements

The authors wish to acknowledge support from Beijing University of Chinese to Yuanyuan Shi (No.1000061020086).

## DATA AVAILABILITY

The mass spectrometry proteomics data have been deposited to the iProX integrated proteome resources via the iProX web (https://www.iprox.org/), IPX ID: IPX0002335000.

---

Supplementary Figure 1. A. Overlap of up-regulated proteins in model group and down-regulated proteins in other groups. B. Restored proteins in ESDH for arthritis model. C. Volcanic map of changed proteins in arthritis *vs*. control. D. mRNA expression of OA changed genes. Data are expressed as mean ± SD (D, pooling samples, *n*=3).

Supplementary Figure 2. The original western blot results for the experiments.

## Supplementary Table legends

Supplementary Table 1 LC-MS/MS for the identification of components in ESD

Supplementary Table 2 Identified proteins through LC-MS/MS analysis

Supplementary Table 3 Quantified proteins through label-free proteomics analysis

Supplementary Table 4 Quantified proteins through label-free proteomics analysis (unique piptides>=2)

Supplementary Table 5 Up-regulated proteins in OA model group compared with control

Supplementary Table 6 Down-regulated proteins in OA model group compared with control

Supplementary Table 7 Up-regulated proteins in GA group compared with OA model

Supplementary Table 8 Down-regulated proteins in GA group compared with OA model

Supplementary Table 9 Up-regulated proteins in ESDL group compared with OA model

Supplementary Table 10 Down-regulated proteins in ESDL group compared with OA model

Supplementary Table 11 Up-regulated proteins in ESDH group compared with OA model

Supplementary Table 12 Down-regulated proteins in ESDH group compared with OA model

Supplementary Table 13 HCL clustering results

Supplementary Table 14 Primers list of RT-qPCR

